# Activity-dependent COX-2 proteolysis generates a catalytically inactive fragment that affects aerobic respiration and proliferation

**DOI:** 10.1101/2024.04.08.588559

**Authors:** Liat Hartal Benishay, Sharon Tal, Amal Abd Elkader, Omar Ehsainieh, Ranin Srouji-Eid, Martin Mikl, Liza Barki-Harrington

## Abstract

Cyclooxygenase-2 (COX-2) catalyzes arachidonic acid (AA) into PGH_2_, the single source of all prostaglandins (PGs), ligands that activate multiple inflammatory pathways. AA catalysis quickly results in suicide inactivation, rendering the enzyme catalytically inactive. We show that the catalytic activity also leads to controlled cleavage of COX-2, an event that is differentially regulated by fatty acids, and blocked by COX inhibitors. We also observe COX-2 cleavage in human colon tumors. Using mass spectrometry, we identify two adjacent cleavage points within the catalytic domain, which give rise to COX-2 fragments that are catalytically inactive and localize to different cellular compartments. One of these fragments significantly alters the expression of mitochondrial electron transport genes and functional assays show that it leads to reduced mitochondrial function, increased lactate production, and enhanced proliferation. We propose that in addition to its role in generating PGs, COX-2 has subsequent PG-independent cellular functions that may account for the complex role of COX-2 in proliferative diseases and chronic inflammation.

**Graphical Abstract:** 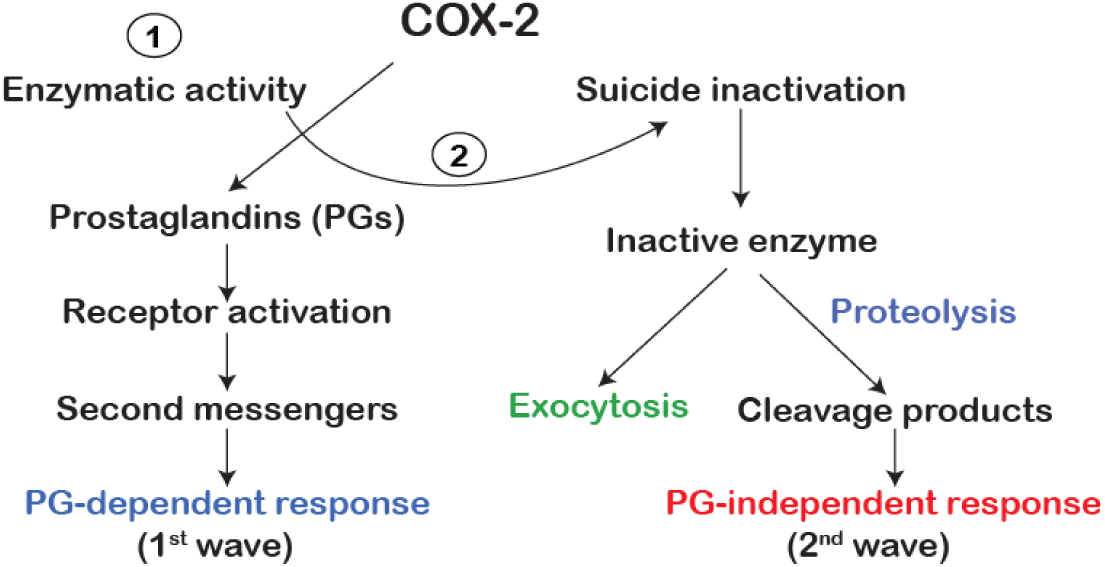

## Introduction

Cyclooxygenase-2 (COX-2) is an enzyme that catalyzes the rate-limiting step in the conversion of arachidonic acid (AA) to prostaglandins (PGs) – bioactive lipids that play central roles in inflammation. Except for several tissues where it is constitutively expressed (*e.g.* kidney, brain), COX-2 expression is usually low, and its levels are rapidly upregulated in response to a wide range of inflammatory and pathological signals, thus producing the PGs that mediate a major part of inflammation ^1^.

COX-2 converts AA into prostaglandin H_2_ (PGH_2_), the source of all subsequent PGs, by a two-step redox reaction, which occurs in the catalytic domain of the protein ^2,3^. The COX-2 enzyme is an obligatory dimer, consisting of two identical monomers, each containing the same active site for AA catalysis. However, only one of the two active sites catalyzes AA ^4,5^, while the other serves as an allosteric modulator of the active site. The rate of catalysis is affected by fatty acids, non-steroidal anti-inflammatory drugs (NSAIDs), and selective COX-2 inhibitors that either compete with AA in the catalytic site or modulate its activity by binding to the allosteric site ^6–9^.

Free radicals formed during catalysis cause COX-2 to quickly become catalytically inactive, by a mechanism of substrate-dependent suicide inactivation ^2,3,10^. Interestingly, whereas the cellular proteasome regularly degrades the quiescent COX-2 that has not been exposed to AA, the fate of the catalytically inactive protein is largely unknown ^11,12^. We have previously shown that some of the substrate-inactivated COX-2 is discarded from the cells by exosomes ^13^. Now we show that AA-stimulation also causes the COX-2 protein to undergo limited proteolysis and that one of the resulting fragments inhibits aerobic respiration in a PG-independent manner.

## Results

### A fragment of COX-2 appears following its stimulation with AA

We have previously shown that a brief stimulation of HEK 293 cells overexpressing COX-2 with AA, causes the appearance of a smaller COX-2 immunoreactive band of ~40 kD ^13^. To determine whether this is a general phenotype of COX-2, we stimulated multiple human cancer-derived cell lines with endogenous COX-2 expression, with AA, and probed the samples with an anti-COX-2 antibody directed against the N-terminal of the protein. As depicted in Fig. 1A, AA stimulation caused the appearance of a ~40 KD COX-2 immunoreactive band in all cell lines examined. Some cells (e.g. A549, HeLa, MCF10A) also expressed additional COX-2 immunoreactive bands that were less affected by the presence of AA. An immunoreactive band of the same size was also observed after AA stimulation in the murine-derived RAW 264.7 macrophages (Fig. 1B) and in HEK 293 cells with exogenous expression of COX-2 (Fig. 1C). Together, these data suggest that the AA-dependent appearance of the lower COX-2 immunoreactive band is a general phenomenon that is not restricted to a certain cell line or species or to whether COX-2 is endogenously or ectopically-expressed.

**Fig. 1:**
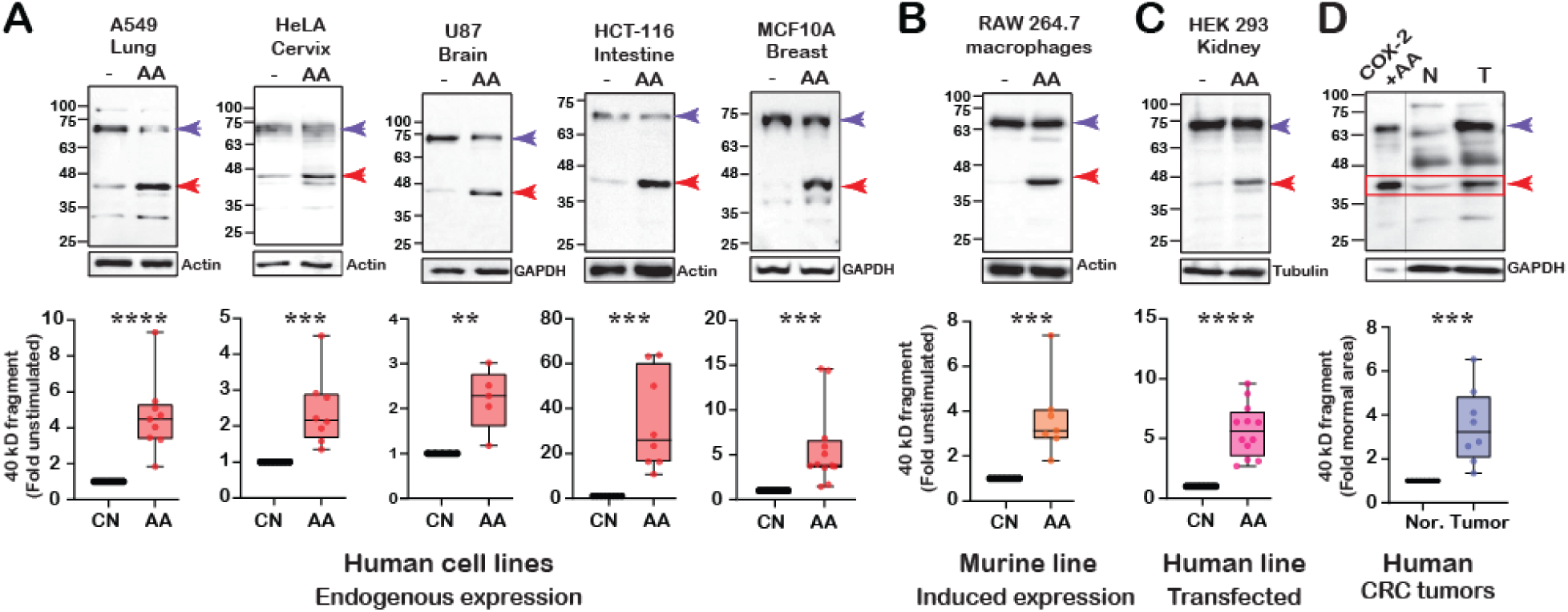
A fragment of COX-2 appears following its stimulation with AA. (A) Representative western blots (upper panels) and quantification (lower panels) of data from five human cancer-derived cell lines with endogenous COX-2 expression, before and after stimulation with arachidonic acid (AA). A549 (n=9), HeLa (n=8), U87 (n=5), HCT-116 (n=8), and MCF10A (n=11). (B) Representative western blots (upper panel) and quantification (lower panel) of data from murine RAW 264.7 cells, with overnight LPS-induced COX-2 expression, before and after stimulation with AA (n=6). (C) Representative western blots (upper panel) and quantification (lower panel) of data from COX-2 transfected HEK 293 cells, before and after stimulation with AA (n=12). (D) Representative samples from a tumor (T) and surrounding normal tissues (N) obtained from a colorectal cancer patient, ran alongside a sample of COX-2 expressing HEK 293 cells stimulated with AA, and quantification of n=8 samples. Statistical analysis was performed using the Wilcoxon test. Quantification of the protein levels of the COX-2 fragment was done after normalization by GAPDH or actin. Data are represented as mean ± SD. Statistical significance was assessed by Student’s t-test. ^∗^p < 0.05, ^∗∗^p < 0.01, ^∗∗∗^p < 0.001, and ^∗∗∗∗^p < 0.0001.

We next examined the expression of COX-2 in a small cohort of eight samples of colorectal tumors of patients diagnosed with low or moderate colorectal adenocarcinoma ^14^. As depicted in Fig. 1D, cancerous samples also showed elevated expression of a ~40 kD immunoreactive band compared to tissue obtained from the clean margins of the same sample. This fragment was the same size as the one that appeared in AA-stimulated COX-2 expressing cells (Fig. 1D, lane 1). While the sample size of the patient cohort is too small to make any meaningful correlations (e.g. pathology, severity, prognosis, etc.), these findings suggest that smaller COX-2 fragments exist in vivo, and hence support the rationale for searching for their identity and possible function in the cell.

### The appearance of the COX-2 fragment requires enzymatic activity

Following binding to COX-2, AA is catalyzed in two structurally and functionally interconnected sites within the protein. The first is a cyclooxygenase (COX) reaction, which takes place in a hydrophobic channel in the core of the enzyme where prostaglandin endoperoxide G_2_ (PGG_2_) is generated. The subsequent peroxidase (POX) reaction occurs at a heme-containing active site located near the protein surface where PGG_2_ is reduced to PGH_2_. Specific PG synthases then catalyze PGH_2_ into PGs (PGE_2_, PGD_2,_ PGI_2,_ PGF_1α_) and thromboxane (TXA_2_), all of which activate multiple signaling pathways by binding to G protein-coupled receptors ^2,15^. To determine which of these stages is necessary for AA-mediated generation of the COX-2 fragment, we first tested whether the final products of AA catalysis can cause the appearance of this band. For this, we treated COX-2-expressing cells with PGE_2_, one of the most abundant products of AA catalysis, for 30-120 min. As depicted in Fig. 2A, in contrast to AA stimulation, PGE_2_ treatment did not cause the appearance of the COX-2 fragment at any time, suggesting that this phenomenon is probably not mediated by the soluble products of AA catalysis.

**Fig. 2:**
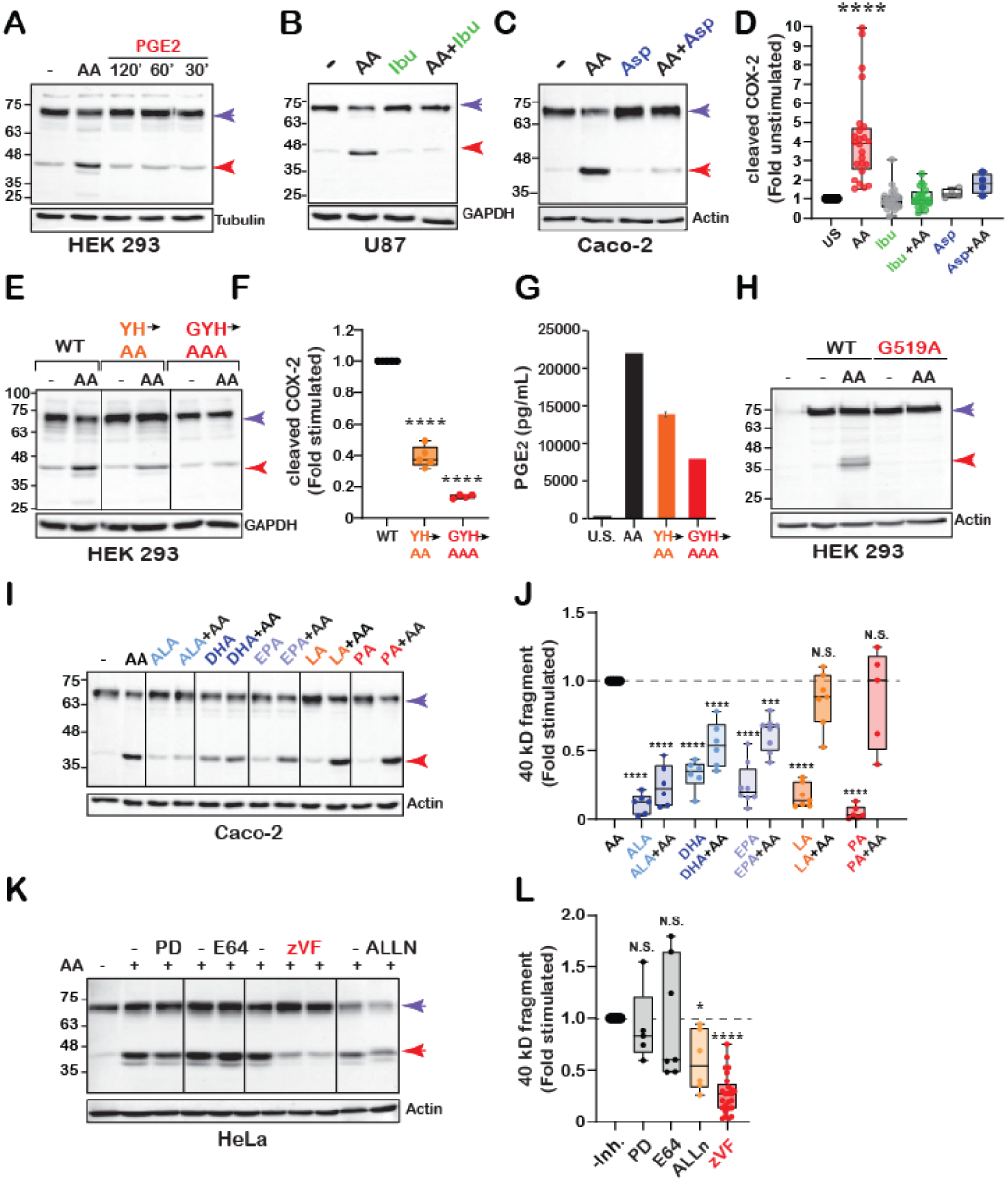
The appearance of the COX-2 fragment requires enzymatic activity. (A) Western blot of COX-2 expressing HEK 293 cells, treated with either AA or PGE_2_ at different time points. (B) Western blot of U87 cells stimulated with AA in the absence or presence of ibuprofen (Ibu). (C) Western blot of Caco-2 cells stimulated with AA in the absence or presence of aspirin (Asp). (D) Box and whisker plot illustrating the quantification of average changes in the levels of the COX-2 fragment in response to ibuprofen. A total of n=24 consisting of data obtained from U87 (n=4), A549 (n=6), HeLa (n=2), RAW 264.7 (n=3), Caco-2 (n=5), HCT-116 (n=4) for ibuprofen or aspirin (Caco-2, n=4). (E) Western blot depicting the response of HEK 293 cells expressing either WT, double mutation (Y341A/H342A, orange), or triple mutation (G340A/Y341A/H342A) COX-2 to AA stimulation. (F) Quantification of the COX-2 fragment levels in (E) after normalization by GAPDH (n=4). (G) Quantification of PGE_2_ levels produced by stimulation of HEK 293 cells expressing WT and COX-2 mutants (n=2). (H) Representative western blot of Caco-2 cells treated with different fatty acids (100 μM each) 30 min before AA stimulation. (I) Quantification of COX-2 fragment levels in (I) after normalization by actin. ALA (n=6), DHA (n=6), EPA (n=8), LA (n=7), PA (n=5). (J) Representative blots of HeLa cells treated with the protease inhibitors PD150606 (PD, 50 μM), E64D (10 μM), zVF (50 and 100 μM), and ALLN (10 μM) overnight, before AA stimulation. (K) Quantification of COX-2 fragment levels in (K) after normalization by GAPDH or actin in response following treatment with protease inhibitors. PD150606 (n=5), E64 (n=7), ALLN (n=6), zVF (n=25). Statistical analysis was assessed by One-way ANOVA ^∗∗∗∗^p < 0.0001, with Dunnett’s multiple comparisons post-test.

Since the appearance of the COX-2 fragment was dependent upon the presence of AA, we next pretreated the cells with reversible or irreversible inhibitors of COX-2 (ibuprofen and aspirin, respectively) before stimulation with AA. The results presented in Fig. 2B-D show that treatment with both compounds blocked the appearance of the COX-2 fragment. Ibuprofen and aspirin both inhibit catalytic activity by preventing access of AA to the catalytic sites ^2^. We therefore postulated that the generation of the COX-2 fragment requires either binding and/or catalytic processing of AA. A productive binding mode of AA to the enzyme is obtained when it is positioned such that the carboxylate lies near amino acids Y106 and Y341, at the opening of the cyclooxygenase channel ^16^. We therefore introduced point mutations to Y341 and its vicinity and explored the effect on the formation of the COX-2 fragment. As depicted in Fig. 2E-F, a double mutation in Y341A/H342A caused a marked reduction in the appearance of the fragment in response to AA, and an additional mutation at G340 (G340A/Y341A/H342A) prevented it altogether. These mutations also caused a severe reduction in the catalytic activity of COX-2 as measured by its ability to generate PGE_2_ (Fig. 2G). Finally, a COX-2 mutant that binds AA but shows minimal enzymatic activity compared to the native enzyme (G519A) ^15^, also prevented the appearance of this band (Fig. 2H). Together, these data indicate that AA-mediated formation of the COX-2 fragment requires both productive binding and catalytic processing of AA, as attenuation of AA binding and impairment of catalysis either chemically or genetically prevents its formation.

The rate of AA catalysis by COX-2 was previously shown to be differentially modulated by the binding of both substrate- and non-substrate fatty acids (FAs) ^4,6,9^. To determine if FAs also affect the appearance of the COX-2 fragment, we treated COX-2-expressing cells with saturated and unsaturated (ω3 and ω6) FAs before AA stimulation. As depicted in Fig. 2I-J, the ω3 FAs dihomo-γ linoleic acid (DHA, 22:6ω3), α linolenic acid (ALA, 18:3ω3), and eicosapentaenoic acid (EPA, 20:5ω3) all caused a slight increase in the formation of the fragment when applied alone. Remarkably, when applied before stimulation with AA, all three FAs severely reduced the appearance of the COX-2 fragment. In contrast, linoleic acid (LA, 18:2ω6) or palmitic acid (PA, 16:0) did not affect its appearance following activation. These findings suggest that the formation of the COX-2 fragment following enzymatic activity is a regulated event, which is affected by the content of FAs present.

Our data so far indicates that the appearance of the COX-2 fragment is a post-translational event that may be carried out by proteolysis. Previous studies had indicated that cleavage of the full-length (F.L.) COX-2 is reduced by different cysteine protease inhibitors, among which calpain inhibitors had the strongest effect ^17,18^. To test whether calpain, a calcium-dependent cysteine protease is also involved in AA-mediated generation of the COX-2 fragment, we tested its formation in the presence of an array of general cysteine protease inhibitors and selective calpain inhibitors. As shown in Fig. 2K, E64 (a broad-spectrum cysteine protease inhibitor) ^18^ and PD150606 (a selective calpain inhibitor) ^19^ did not affect the AA-mediated formation of the COX-2 fragment. Application of ALLN (calpain inhibitor I) reduced the appearance of the fragment by approximately 60 %, and Z-Val Phe-OH (zVF, Calpain III), a potent cell-permeable calpain I and II, had the most profound effect, inhibiting approximately 75 % of cleavage (Fig. 2L).

### Identification and ectopic expression of COX-2 fragments

Given the above findings, we next sought to identify AA-mediated cleavage points in COX-2. For this purpose, we stimulated RAW 264.7 cells with LPS to induce COX-2 expression, followed by exposure to AA to induce cleavage. Samples were then separated by SDS-PAGE, and proteins were extracted and subjected to Liquid Chromatography with tandem mass spectrometry (LC-MS/MS) analysis. Total lysates from the same experiments were analyzed by western blot in parallel, to confirm the appearance of the fragment. Probing the membranes with anti-COX-2 against the N-terminal of COX-2 identified the F.L. COX-2 as well as the additional ~40 kD band that was detected in the other cell lines (Fig. 3A and Fig. 1). The use of an additional antibody directed against the C-terminal of COX-2 detected the F.L. form and a prominent band of ~ 30 kD (Fig. 3B). Given that the size of F.L. COX-2 is ~72 kD, these results suggest that at least one cleavage point is present roughly in the middle of the protein, one which gives rise to two complementary fragments of ~40 and ~30 kD.

**Fig. 3:**
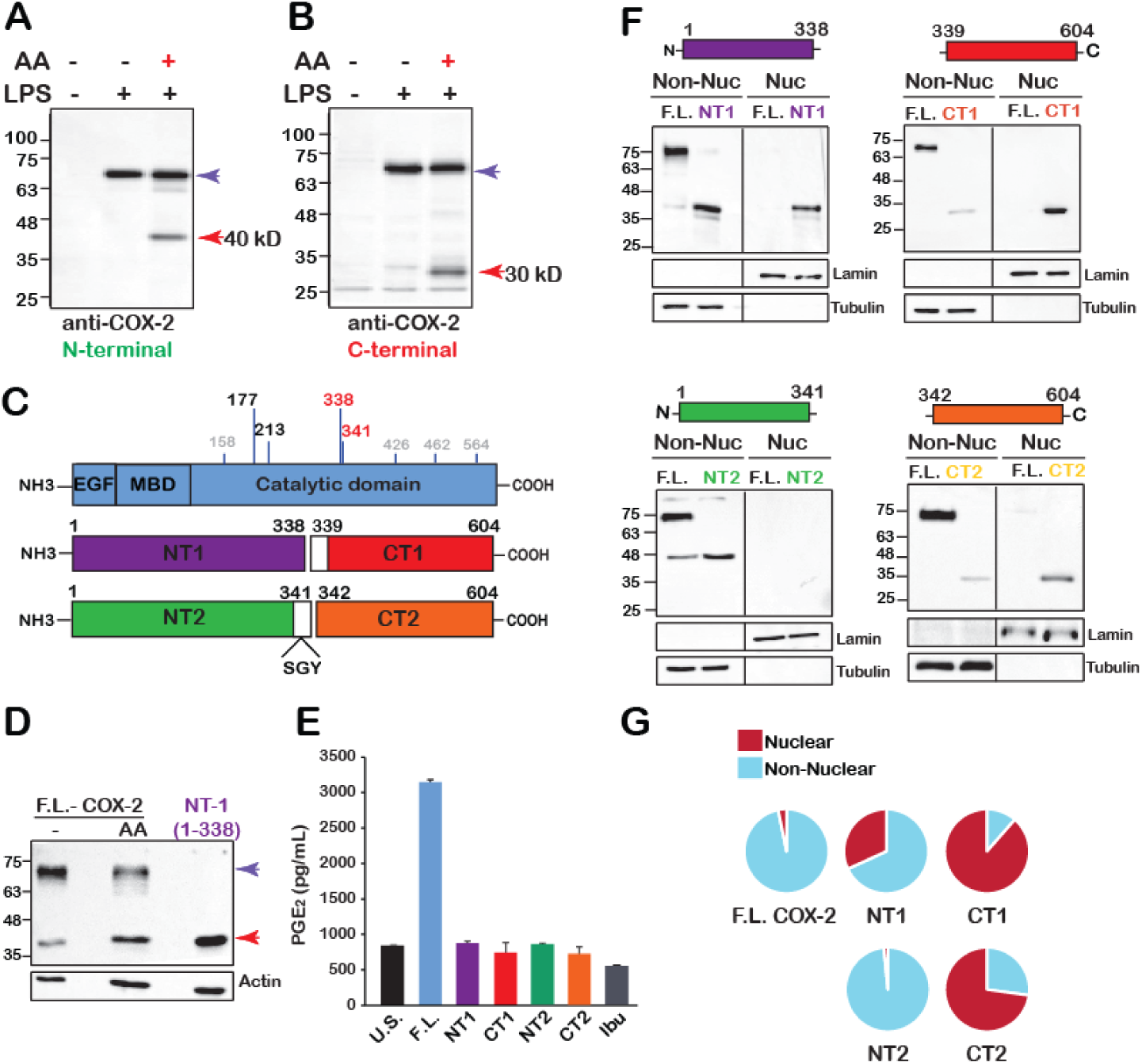
Identification and ectopic expression of COX-2 fragments. (A-B) Western blots of RAW 264.7 cells stimulated overnight with LPS before exposure to AA using antibodies against the amino- (A) and carboxy- (B) termini of COX-2. (C) Schematic depiction of the eight cleavage sites identified by LC-MS/MS in RAW 264.7 cells and the overlapping amino acids in homo sapiens (Top drawing). Schematic representations of the putative cleavage products that were introduced into cells. NT1=1-338, CT1=339-604 (middle drawing), and NT2=1-341 and CT2=342-604 (bottom drawing). (D) Western blot depicting NT1 relative to AA-stimulated COX-2 in HEK 293 cells. (E) Quantification of PGE_2_ levels produced by HEK 293 cells expressing WT COX-2 or the different fragments. U.S. represents cells expressing WT COX-2 without stimulation, and Ibu represents cells treated with ibuprofen for 30 min before AA stimulation, (n=2). (F) Representative immunoblots of nuclear (Nuc) and non-nuclear (Non-Nuc) fractions of HEK293 cells expressing F.L. COX-2 or each of the four fragments. (G) Pie charts quantifications of COX-2 or fragments after normalization of nuclear fractions to lamin B and non-nuclear fractions to tubulin. NT1 (n=7), CT1 (n=5), NT2 (n=4), CT2 (n=7).

Analysis of three independent LC-MS/MS experiments yielded eight putative cleavage points, four of which were positioned at amino acids that are identical to those of human COX-2 (177, 213, 338, and 341, Fig. 3C). Interestingly, the estimated mass of two cleavage products that were very close to one another (positions 338-339 and 341-342, marked in red) were ~40 kD (1-338/1-341) and 30 kD (339-604 and 342-604) respectively, which is similar to the estimated mass of the fragments obtained by western blot (Fig. 3A-B). To confirm this, we ectopically expressed one of the putative fragments (1-338, NT1 herein) in HEK 293 cells, and ran the sample alongside a sample of AA-stimulated COX-2. As shown in Fig. 3D, NT1 migrated to the same distance as the 40 kD COX-2 immunoreactive fragment that appeared following AA stimulation.

Our data show that cleavage of COX-2 is a post-translational event that must be preceded by enzymatic activity (Fig. 2). Therefore, studying the possible effects of the AA-dependent fragments cannot be done simply by activation of COX-2 with AA, since this will yield a mixed response of PG-dependent and independent effects. To separate between the two, we cloned the sequences of two pairs of putative COX-2 fragments that complement each other to the size of F.L. COX-2: NT1 (amino acids, aa 1-338, Fig. 3D) and its complementary CT1 (aa 339-604) and NT2 (aa 1-341) and its complementary CT2 (aa 342-604), and expressed each one ectopically. We then confirmed that in contrast to F.L. COX-2, none of these fragments can generate PGE_2,_ indicating that they lack enzymatic activity (Fig. 3E).

The COX-2 homodimer is associated with the ER and nuclear membranes by four amphipathic helices of its membrane binding domain (MBD). These chains surround the opening of the cyclooxygenase channel through which AA enters the COX site ^20^. Since amino acids 338-342 are located near the opening of the cyclooxygenase channel ^16^, we hypothesized that loss of the MBD motif due to cleavage may result in changes in the cellular localization of some of the fragments, particularly CT 1 and CT2, compared to that of the F.L. protein. To test this, we examined the cellular localization of the F.L. COX-2 and each of the four fragments. As depicted in Fig. 3F-G, F.L. COX-2 appears almost entirely in the non-nuclear fractions (97%) as indicated by staining with antibodies against both the N- and C-termini (left lane of all images in Fig. 3F). The same is true for fragment NT2 (99%) but not for NT1, 32% of which is found in nuclear fractions (Fig. 3F top left panel and Fig. 3G). In marked contrast to F.L. and NT fragments, both CT1 and CT2 localize primarily to the nuclear fractions, with CT1 localizing almost completely to that fraction (90% and 73%, respectively, Fig. 3F-G).

### CT1-COX-2 reduces mitochondrial function and increases lactate production

The marked differences in localization of CT1 and CT2 compared to F.L. COX-2 led us to postulate that they may differentially affect cellular pathways by altering gene expression. To test this hypothesis, we sequenced the transcriptomes of control (untransfected, UT) and cells with stable expression of CT1 or CT2. A principal component analysis (PCA) of the resulting gene-level expression measurements revealed that both CT1 and CT2 exhibit different transcriptomic profiles compared to the control cells as well as to each other (Fig. 4A). Analysis of the RNA-seq data identified a total of 1852 genes that were differentially expressed (DE) between CT1 and CT2, (1008 genes, 54% upregulated (Table S1) and 844, 46% downregulated (Table S2); Benjamini-Hochberg adjusted *p*<0.05) (Fig. 4B), even though the two CT fragments differ only in the additional three N-terminal amino acids of CT1. To select for effects that were specific to the longer CT1 fragment that was almost exclusively found in nuclear fractions, we performed an additional filtering step for genes that also changed between CT2 and UT cells, which resulted in a list of 608 genes that were uniquely affected by the expression of CT1, 295 (49%) of which were upregulated (Table S3) and 313 (51%) were downregulated (Benjamini-Hochberg adjusted p<0.05) (Tables S4).

**Fig. 4:**
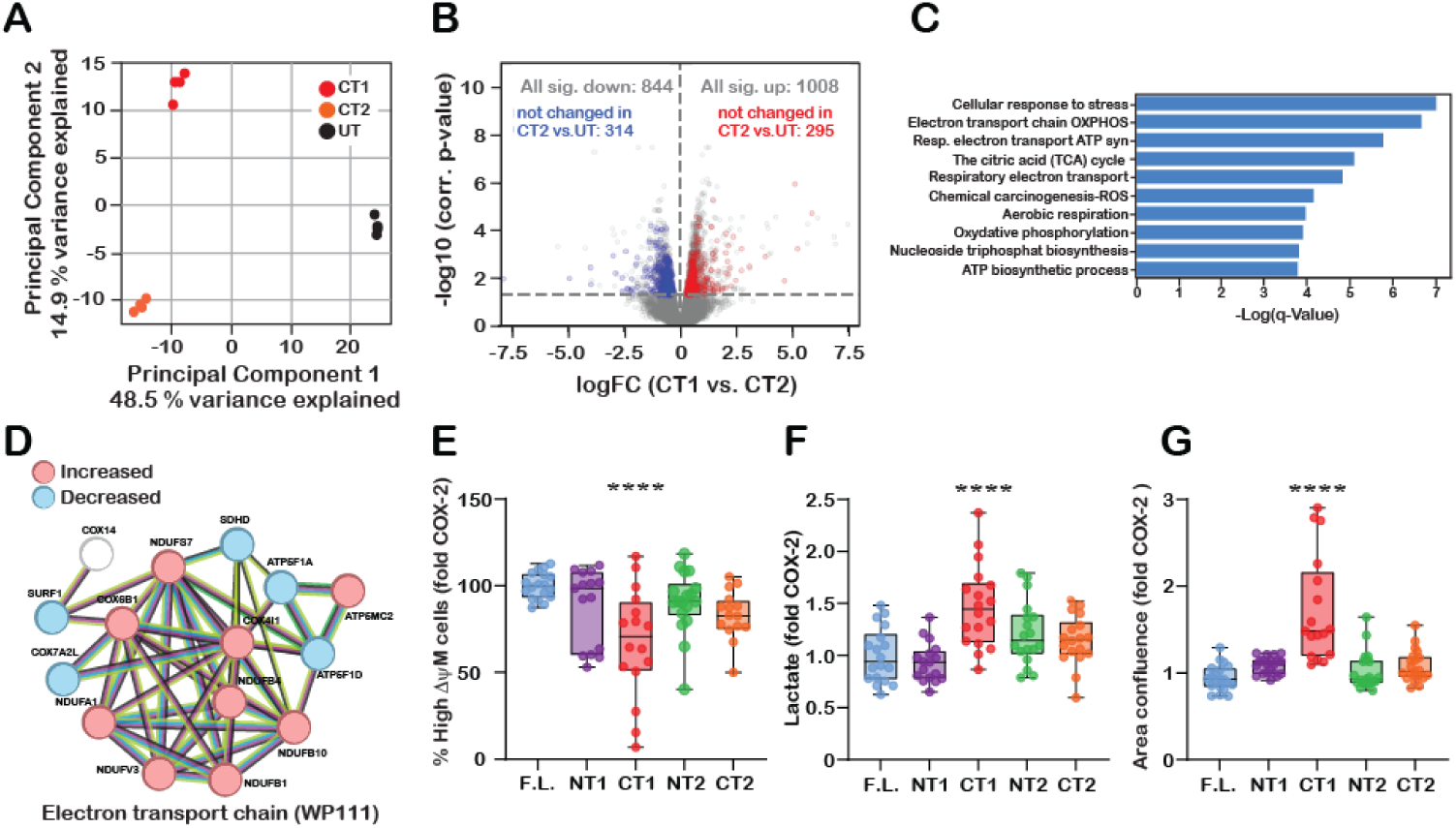
CT1-COX-2 reduces mitochondrial function and increases lactate production. (A) Scatter plot showing the first two principal components from a PCA based on normalized and standardized expression values of all genes covered by at least 10 reads in all samples. (B) Volcano plot depicting the differential expression of genes between CT1- and CT2-expressing cells. The horizontal dashed line indicates the significance cutoff (adjusted p-value<0.05). Red and blue data points indicate genes that change between CT1- and CT2-expressing cells but do not exhibit a significant increase or decrease in CT2-expressing compared to control cells. (C) Top 10 most significantly enriched functional groups (as determined using Metascape) among the upregulated genes in CT1-compared to CT2-expressing cells, which showed no significant change between CT2-expressing and control cells. (D) STRING analysis of a DE gene set showing a minimum interaction score of highest confidence (0.9). (E) Box and whiskers plot describing the effect of COX-2 fragments on mitochondrial function, normalized to WT COX-2: F.L. COX-2 (n=19), NT1 (n=15), CT1 (n=16), NT2 (n=16), CT2 (n=15). (F) Box and whiskers plot describing the effect of COX-2 fragments on lactate production, normalized to WT COX-2: F.L. COX-2 (n=18), NT1 (n=17), CT1 (n=18), NT2 (n=18) CT2 (n=18). (G) Box and whiskers plot describing the effect of COX-2 fragments on proliferation, normalized to WT COX-2: F.L. COX-2 (n=20), NT1 (n=20), CT1 (n=17), NT2 (n=21) CT2 (n=21). One-Way ANOVA, *** p<0.001 vs. COX-2 (Dunnett’s multiple comparisons post-test).

We next used Metascape ^21^ to identify the pathways that are enriched among the DE genes (Tables S3 and S4). This analysis revealed a significant enrichment of genes belonging to cellular aerobic respiration among the genes upregulated in CT1-expressing cells (e.g. Electron transport chain OXPHOS system in mitochondria, WP111, q = 2.0*10^-7^; Respiratory Electron Transport, R-HAS-163200, q = 1.5*10^-6^; aerobic respiration, GO: 0009060, q = 1.1*10^-4^) (Fig. 4C, Table S5). The genes downregulated in CT1-expressing cells showed less significant enrichment for functional groups (Figure S1, Table S6). Application of the STRING database ^22^ on all DE genes revealed a network of 17 genes belonging to the electron transport chain (WP111 Electron transport chain OXPHOS system in mitochondria, Fig. 4D). Interestingly, the effect on the electron transport chain was mixed: genes belonging to complex I (NADH-Ubiquinone oxidoreductase) were all elevated, whereas those belonging to complex II and V (Succinate-Ubiquinone oxidoreductase, and ATP synthase, respectively) were downregulated. Genes belonging to complex IV showed a mixed effect where some genes were upregulated while others were downregulated (Fig. 4D).

Based on the results of the RNA-seq analysis, we next measured the effect of all COX-2 fragments on mitochondrial membrane potential using tetramethylrhodamine ethyl ester (TMRE), as described ^23^. Remarkably, only cells that expressed CT1 presented a significant reduction in the levels of high-membrane potential mitochondria (Fig. 4E). Reduced aerobic respiration can potentially lead to the generation of ATP by the anaerobic glycolysis pathway, which leads to an increase in lactate production. We therefore measured lactate levels in the supernatant of cells expressing the four fragments (Fig 4F). The data depicted in Fig. 4F confirmed that compared to cells expressing F.L. COX-2 only cells that expressed CT1 exhibited elevated lactate levels. A reduction in the activity of oxidative phosphorylation despite the presence of oxygen is one of the hallmarks of rewiring of cancer cell metabolism. This phenomenon termed the Warburg effect, is characterized by a significant increase in lactate production due to the conversion of pyruvate to lactate instead of its metabolism by the mitochondria ^24^. Given the effect of CT1 on mitochondrial function and lactate production, we next measured the effect of COX-2 and its fragments on the rate of cell proliferation. As depicted in Fig. 4G, the only COX-2 fragment that caused a consistent and significant increase in proliferation rate was CT1.

## Discussion

The main finding of the present study is that following AA-induced catalytic activity, COX-2 undergoes limited proteolysis, and at least one of the resulting fragments affects aerobic respiration and rate of proliferation. Hence, we propose that COX-2 fulfills two different roles in the cell. The first is its undisputed role as a central enzyme that generates soluble ligands of inflammation, and the second is a PG-independent role that is mediated through interactions of COX-2 fragments with different cellular pathways.

The structure and function of the COX-2 enzyme have been studied in-depth, providing detailed information regarding the kinetics of AA catalysis, and interactions of FAs and COX inhibitors with its catalytic and allosteric sites ^6,7,9,15^. In terms of its expression, COX-2 is an immediate-early gene that is rapidly and massively upregulated to meet the high demand for PGs during inflammation, and elevated COX-2 levels are hallmarks of chronic inflammation and many types of malignancies ^25^. Therefore, the majority of the high-volume literature regarding COX-2 focuses on its transcriptional regulation ^1^ and on pharmacological means to inhibit its enzymatic activity ^26^. In contrast, much less is known about pathways that clear COX-2 from the cell. Previous studies have indicated that in the absence of AA, COX-2 is degraded in the cellular proteasome, but that the activated protein is no longer degraded there ^11,12^. This is a very important matter that is largely overlooked, especially since one of the most important properties of COX-2 is that its ability to generate PGs is limited by self-induced suicide inactivation occurring within seconds of activation ^27^. Previously, we demonstrated that part of the substrate-inactivated protein is discarded from the cells by exosomes ^13^, and we now further bridge the gap by showing that COX-2 is also subjected to controlled cleavage, which generates fragments that have a secondary effect on cells.

COX-2 was previously shown to undergo proteolysis into discrete fragments following exposure of human synovial fibroblasts to pro-inflammatory cytokines ^18^. Using an anti-pan-human COX-2 with an undisclosed epitope, the researchers identified four COX-2 fragments, two of which are similar to those identified by us (44-42 and 34-36 kD). Using the same antibody against the C-terminus of COX-2 as we have (Fig. 3B), they also identified a fragment of 30 kD, which like CT1 and CT2 (Fig. 3G), localizes to nuclear and mitochondrial-lysosomal fractions. The similarity between the sizes of fragments in both studies is probably because the cytokine IL-1β causes AA release ^28^, which is comparable to the application of AA directly onto the cells. However, while Mancini et al ascribe a role for proteolysis in protein maturation, we show that the putative fragments completely lack enzymatic activity, thus suggesting that they affect the cell in a PG-independent manner. These results are supported by our previously published finding that COX-2 which lacks the EGF and MBD domains (ΔCOX-2) is catalytically inactive ^29^.

The search for protease/s that cleave COX-2 is challenging. Whereas the F.L. unstimulated COX-2 is degraded in the proteasome, it no longer does so after catalytic activity ^12^. Since the putative cleavage site in our study lies at the surface of the protein ^16^, and given earlier indications ^17,18^, we examined calpain as a possible candidate for COX-2 proteolysis. Calpain was shown to degrade COX-2 in vitro ^17^, and indeed calpain inhibitors such as the calpain III inhibitor zVF significantly decreased the appearance of the 40 kD fragment (Fig. 2K-L). However, zVF also targets γ-secretase ^30^, and even though it inhibited the majority of cleavage, it did not obliterate it, even in the presence of a cycloheximide that inhibits *de novo* synthesis (not shown). Therefore, the identity of the protease/s that cleave COX-2 remains to be determined.

Multiple lines of evidence, both in cell lines and *in vivo*, indicate that COX-2 is involved in the initiation, promotion, and progression of cancer ^31,32^, and many studies show that elevated expression of COX-2 is strongly associated with a poor prognostic outcome ^33–35^. The role of COX-2 in tumorigenesis is attributed to the production of PGs that enhance proliferation by stimulation of their respective receptors ^36,37^ thereby activating downstream signaling pathways that are associated with regulation of the cell cycle and proliferation ^38–39^. While there is a general agreement that inhibition of COX-2 is advantageous in cancer prevention ^40–46^, the efficiency of these drugs after disease onset is less straightforward. Several comprehensive studies showed that combining chemotherapy with COX-2 inhibitors in treating human cancers, is significantly beneficial in reducing tumor burden ^31,47^, while other large-scale ones found that celecoxib-combined therapy failed to improve disease-free survival or progression-free survival, and had no effect on pathological complete response ^48^. Clinical trials in malignancies where COX-2 expression correlates with poor prognosis (e.g. lung, breast, glioblastoma, pancreas, and colon) failed to show significant advantages for the use of NSAIDs or Coxibs in disease treatment ^49–54^. Without contradicting the proven role of the PG-mediated role of COX-2 in proliferative diseases, we propose that the AA-induced COX-2 cleavage products present an additional non-catalytic aspect of COX-2 signaling that can modulate (exacerbate, or mitigate) the initiation and/or course of proliferation and should be explored. We also suggest that the presence of COX-2 fragments in tumors is not detected because the primary method of detection in the clinical setting is immunohistochemistry, which is excellent for determining the amount of expression and tissue localization but does not indicate the size of the protein (e.g. Fig. 1D).

A non-catalytic role has been ascribed to COX-2 in a series of studies showing that it inhibits the function of the tumor suppressor p53 through a direct interaction between the two proteins ^55–57^. In contrast to our findings, the effect of COX-2 on p53 was found to occur without stimulation. Furthermore, p53 interacts with the non-catalytic regions of COX-2 (aa 1-126), while we identify effects that are mediated by regions of the catalytic domain of the protein (aa 339-604). Nonetheless, both studies complement each other by supporting additional cellular functions for COX-2, beyond those obtained by the generation of PGs

In summary, we propose that COX-2 fulfills two different cellular functions that occur in two waves. The first wave is through its enzymatic action and the generation of PGs that activate G protein-coupled receptors and relay their signals through soluble second messengers. The second wave arises from the secession of catalytic activity, which leads to the secretion of part of the inactive F.L. protein by exocytosis ^13^ as well as to its controlled proteolysis. The products of proteolysis can then migrate to different cellular compartments where they affect cellular function in mechanisms that are PG-independent. Characterization of these pathways and discovery of additional functions of COX-2 fragments will further our understanding of the complex role of COX-2 in pathologies such as proliferative diseases and chronic inflammation and may help identify novel targets for intervention against its components.

## Acknowledgments

The authors wish to thank Dr. Sagie Schif-Zuck from the Flow Cytometry Service Unit, University of Haifa for assistance with the flow cytometry analyses, and Dr. Maya Lalzar from the Bioinformatics Services Unit, University of Haifa, for assistance with transcriptome data. This work was supported by the Israel Science Foundation personal grant (1445/14 and 2240/19), and by the Israel Cancer Association (#20210063) to L. B-H.

## Author contributions

Conceptualization, S.T and L.B-H; Methodology, S.T., and L.B-H; Investigation, L.H-B., S.T., R.S-E, A.A., O.E., M.M., L. B-H.; Writing-Original draft, L.B-H; Writing-Review & Editing, L.B-H, M.M.; Funding Acquisition, L. B-H.; Resources, M.M., L.B-H., Supervision, L.B-H.

## Declaration of Interests

The authors declare no competing interests.

## STAR Methods

| REAGENT or RESOURCE | SOURCE | IDENTIFIER |
| --- | --- | --- |
| **Antibodies** |  |  |
| COX-2 Rabbit monoclonal Ab | Cell Signaling Technology | 12282 (H5H5) XP |
| COX-2 (C-20) goat polyclonal Ab | Santa Cruz Biotechnology Inc. | SC-1745 |
| Beta Actin rabbit polyclonal Ab | Abcam | 115777 |
| GAPDH (D4C6R) monoclonal Ab | Cell Signaling Technology |  |
| Lamin B | Cell Signaling Technology | CST-13435 |
| Tubulin | Abcam | 6160 |
| Horseradish peroxidase-conjugated goat anti-rabbit IgG | Jackson ImmunoResearch Laboratories | 111-035-144 |
| Horseradish peroxidase-conjugated goat anti-mouse IgG | Jackson ImmunoResearch Laboratories | 115-035-003 |
| Peroxidase AffiniPure™ Bovine Anti-Goat IgG (H+L) | Jackson ImmunoResearch Laboratories | 805-035-180 |
| **Bacterial and virus strains** |  |  |
| **Biological samples** |  |  |
| Colorectal biopsies (human) | HaEmek Medical Center | Ha'Emek Medical Center- 0049–19 |
| **Chemicals, peptides, and recombinant proteins** |  |  |
| Arachidonic acid | Cayman Chemical | 90010 |
| Aspirin | Cayman Chemical | 70260 |
| Alpha linolenic acid | Cayman Chemical | 90210 |
| Beta-Nicotinamide adenine dinucleotide hydrate | Sigma-Aldrich | N7004 |
| BLUEYE prestained protein ladder | BIO-HELIX | IMPM007 |
| Calpain inhibitor I (ALLN) | Sigma-Aldrich | 11086090001 |
| Calpain inhibitor III -zvf | Sigma-Aldrich | 208722 |
| Complete(TM) Protease Inhibitor Cocktail | Sigma Aldrich | 11697498001 |
| Docosahexaenoic Acid (DHA) | Cayman Chemical | 90310 |
| DAPT | Cayman Chemical | 13197 |
| E64 | Sigma Aldrich | E3132 |
| Eicosapentaenoic Acid (sodium salt) | Cayman Chemical | 21908-5 |
| Hygromycin B | Sigma Aldrich | H3274 |
| Ibuprofen | Cayman Chemical | 16793 |
| Imperial™ Protein Stain Coomassie dye | Rhenium | 24615 |
| Iodonitrotetrazolium chloride | Sigma-Aldrich | I8377 |
| L-lactate dehydrogenase (LDHA) | Sigma-Aldrich | 10127230001 |
| L-lactate (sodium) | Sigma-Aldrich | PHR1113 |
| Lipopolysaccharide (LPS) | Sigma Aldrich | L6529 |
| Linoleic acid | Sigma-Aldrich | L1012 |
| Methoxy-PMS | BioTag | HY-D0937 |
| Palmitic Acid | Cayman Chemical | 10006627 |
| PD 15606 | Sigma Aldrich | D5946 |
| PolyJet In Vitro DNA Transfection Reagent | SingaGen | SL100688 |
| Prostaglandin E2 | Cayman Chemical | 14010 |
| Protein Assay Dye Reagent Concentrate | Bio-Rad | 5000006 |
| Protein A/G PLUS-Agarose | Santa Cruz Biotechnology | SC-2003 |
| RNase-ExitusPlus | Satorius | 01-897-1B |
| TMRE-Mitochondrial Membrane Potential | Abcam | ab113852 |
| Critical commercial assays |  |  |
| Prostaglandin E2 Parameter Assay Kit | R&D Systems, Inc. | KGE004B |
| QuikChange Lightning Site-Directed Mutagenesis Kit | Agilent | 210519 |
| PRESTO™ MINI PLASMID KIT | Hylabs | IMPDH100 |
| Deposited data |  |  |
| Experimental models: Cell lines |  |  |
| A549 | Sheba | Gift Amir Onn |
| Caco-2 | ATCC | HTB-37 |
| HCT-116 | ATCC | CCL-247 |
| HEK 293 | Technion Ins, Israel | Gift Dina Ron |
| HeLa | ATCC | CRM-CCL-2 |
| MCF10A | Technion Ins, Israel | Gift Israel Vlodavsky |
| RAW 264.7 | ATCC | TIB-71 |
| U87 | ATCC | HTB-14 |
| Experimental models: Organisms/strains |  |  |
| Oligonucleotides |  |  |
| Recombinant DNA |  |  |
| Y341A H342A human cox2 (in pcDNA5/FRT/TO) |  |  |
| COX2 G340A Y341A H342A human cox2 (in pcDNA5/FRT/TO) |  |  |
| Software and algorithms |  |  |
| GraphPad Prism 10 |  |  |
| Adobe Illustrator CS5.1 |  |  |
| Adobe Photoshop CS5.1 |  |  |
| ImageQant TL 1D v8.1 |  |  |
| EndNote X7 |  |  |
| Other |  |  |
| Quick-RNA™ Miniprep Plus Kit | Zymo | ZR-R1058 |
| Nitrocellulose blotting membrane- 0.45 µm | Tamar | 980045 |
| PureYield™ Plasmid Midiprep System | Promega | A2492 |
| NEBNext Ultra RNA library Prep kit for Illumina | Thermo Fischer Scientific | E75030L |

### Patient biopsies

Human colon biopsies were obtained from individuals diagnosed and operated on at Ha’Emek Medical Center, Afula Israel. All patients were diagnosed with adenocarcinoma of the colon or rectum, with tumors that were graded at least B2 or higher. Biopsies were obtained during surgery upon written consent from patients according to the Helsinki regulations (Ha’Emek Medical Center-0049–19) and flash-frozen in liquid N_2_ pending analysis. For identification of COX-2 expression patterns were processed exactly using the same samples as we have before ^29^. 150 μg of total tissue lysates were analyzed.

### Cell culture and transfection

A549 lung cancer cells were grown in RPMI. RAW 264.7, U87, HeLa, HEK 293, HCT-116, and Caco-2 were grown in DMEM. All media was supplemented with 10% heat-inactivated fetal bovine serum, 1% pyruvate, 100 μg/mL streptomycin, and 100 U/ml penicillin. MCF-10A cells were grown in DMEM/F12 exactly as described ^58^. Transient transfections were carried out in sub-confluent monolayers (70-80%) using PolyJet (SignaGen Laboratories) at a ratio of 1:3 cDNA: PolyJet, according to the manufacturer’s instructions. HEK293-derived cell lines stably expressing native or fragment COX-2 constructs were generated using hygromycin selection (200 μg/mL medium) for 9 days. The resulting colonies were tested for the presence of each fragment by western blot.

### Cell stimulation, processing, and Immunoblotting

Unless otherwise indicated, all stimulations with AA were at 50 μM for 30 min. All fatty acids used in the study were applied at 100 μM for 30 min before AA stimulation, as were the inhibitors ibuprofen (200 μM) and aspirin (100 μM). Treatments with the different protease inhibitors were applied for 18-24 h at the following concentrations: PD15606 (50 μM), ALLN (10 μM), E64 (10 μM), zVF (50 or 100 μM). Monolayers were washed twice with ice-cold PBS and lysed in RIPA/SDS buffer (50 mM Tris pH-8, 150 mM NaCl, 5 mM EDTA, 1% v/v NP-40, 0.5%, 0.5% w/v deoxycholic acid, 0.1% w/v SDS, 10 mM NaF, 10 mM sodium-pyro-phosphate and Complete Protease Inhibitor cocktail tablets with 0.1 mM PMSF) and process exactly as we have done before ^29^. Subcellular fractionation was done using the Rapid, Efficient And Practical (REAP) method ^59^. The protocol was carried out exactly as described, except for the addition of Complete and 0.1 mM PMSF to the lysis buffer. 40 μg of total protein lysates were used in all blots. Nitrocellulose membranes containing the immuno-complexes or total cell lysate proteins were incubated with primary antibodies at a dilution of 1:500–1000. Proteins were visualized by a WesternBright ECL (Advansta) and quantified using the Amersham Imager 600 (GE) and quantified using ImageQant TL 1D v8.1 software.

### Immunoprecipitation and mass spectrometry

7 x 10^6^ RAW 264.7 cells were grown in 15 cm dishes and stimulated overnight with 100 ng/mL LPS, followed by a 30-minute stimulation with 50 μM AA. The reaction was terminated by two subsequent washes with ice-cold PBS and lysis in 0.5 mL RIPA/SDS buffer. Samples were centrifuged at 14,000 × g at 4 °C for 10 min, the supernatants were collected, and protein levels were determined. 100 μg of the lysates were used for total protein assessment. 3-3.5 mg protein were used for IP experiments. Samples were pre-cleared with 20 μl of protein A/G beads for 60 min, centrifuged again at 14,000 × g at 4 °C for 2 min and the remaining supernatant was used for overnight immunoprecipitation with 2 mg goat polyclonal anti-COX-2 (Santa Cruz Biotechnology, SC-1745) and 30 μl protein A/G beads with continuous rotation at 4 °C. After three subsequent washes with 0.75 mL RIP/SDS, lysates were separated by a 10% sodium dodecyl sulfate (SDS) polyacrylamide gel. The entire protein lane of each sample was cut into three horizontal gel pieces and processed by in-gel trypsin digestion procedure as described before ^60^, followed by peptides desalting using C18 StageTip ^61^. Whole proteome and activity-guided proteasome profiling samples were analyzed using a Q-Exactive Plus mass spectrometer (Thermo Fisher) coupled to Easy nLC 1000 or Dionex (Thermo Fisher). The peptides were resolved by reverse-phase chromatography on 0.075 × 180 mm fused silica capillaries (J&W) packed with Reprosil reversed-phase material (Dr. Maisch; GmbH, Germany). Peptides were eluted with a linear gradient of 6–28% acetonitrile 0.1% formic acid for 60 min followed by a 15 min gradient of 28–95% acetonitrile 0.1% formic acid and a 15 min wash at 95% acetonitrile with 0.1% formic acid in water (at flow rates of 0.15-0.2 μL/min).

The MS analysis was performed in positive mode using a range of m/z 300–1800, resolution of 70,000 for MS1 and 17,500 for MS2, using, repetitively, full MS scan followed by HCD of the 10 most dominant ions selected from the first MS scan with an isolation window of 1.8 m/z. Other settings used were as follows: a dynamic exclusion of 20 seconds, NCE=27, minimum AGC target=8×10^3^, and intensity threshold=1.3×10^5^. Data analysis was done using MSFragger version 4.0 ^62^ via FragPipe version 21.1 (https://fragpipe.nesvilab.org/). In all searches, the raw files were searched against the Uniprot Mouse/Human proteome (Downloaded on Feb 2024 and includes 54940/ 20413 protein sequences, respectively). MSFargger searches were performed by using semi-tryptic digestion. Search criteria included precursor and fragments tolerance of 20 ppm with oxidation of methionine, and protein N-terminal acetylation set as variable modifications.

### COX-2 activity measurements

The media of 3 × 10^5^ transfected HEK 293 in 6-well dishes was siphoned, and the cells were washed twice with warm PBS and incubated in 1 mL of serum-free DMEM with AA for 30min. In the FA experiments, cells were treated with the different FAs (100 μM each) or with the appropriate vehicle for 30 min, followed by an additional 30 min with 50 μM AA. At the end of the experiment, the supernatant was collected and the levels of PGE_2_ were analyzed using the Prostaglandin E_2_ Assay (R&D Systems), according to the manufacturer’s instructions.

### Mitochondrial activity measurements

0.05 x 10^6^ HEK 293 cells were seeded in 12-well dishes and transfected with 0.75 μg F.L. COX-2 or fragment cDNA. 72 h post-transfection the ΔΨm was measured as previously described ^23^. Briefly, the cells were dissociated with trypsin and washed twice with ice-cold 0.2% BSA in PBS. After centrifugation for 5 min at 1500 rpm, the cells were incubated in the dark at 37 °C, for 20 min with Mitochondrial Tracker Green (Ex/Em 490/516) for general mitochondria staining, and TMRE for mitochondrial membrane potential (Ex/Em 549/575 nm). After an additional wash with 1 mL of 0.2% BSA, the cells were resuspended in 300 μl 0.2% BSA and immediately analyzed by the BD FACSCanto II flow cytometer with DACSDiva software (BD Biosciences). Gates were set to exclude necrotic cells and cellular debris and the fluorescence intensity of events within the gated regions was quantified. Data were collected from 10000 events for each sample. The relative percentage of cells with high Δψm was calculated as % high Δψm of total (high Δψm + low Δψm).

### L-Lactate level measurements

L-lactate levels were measured by a colorimetric assay of lactate dehydrogenase (LDH) activity 0.3 x 10^6^ HEK 293 cells were seeded in 6-well dishes and transfected with 1 μg cDNA of F.L. COX-2 or fragments in 1 mL complete DMEM media. 72 h post-transfection the external media of the cells was collected and probed for the presence of lactate exactly according to the step-by-step protocol in https://www.protocols.io/view/colorimetric-determination-of-l-lactate-in-superna-dm6gpj9j5gzp/v1. Briefly, 50 μl of sample supernatants were placed in a flat-bottom 96-well dish (in triplicates) followed by the addition of 50 μl Assay buffer (200 mM Tris-Base, pH=8.2, 2.2 mg/mL β-NAD, 0.5 mg/mL INT, 1 μg/mL L-LDH, 6 μg/mL 1-Methoxy-PMS) and incubated for 1 h at room temperature in the dark. The reaction was terminated by the addition of 50 μl Acetic Acid (1 M) and the absorbance was measured at 490 nm-Ref 650 nm with a microplate absorbance reader. If lactate concentration was above that of the standard curve, samples were diluted in DMEM.

### Proliferation assays

For live cell tracking experiments, 3 × 10^4^ HEK 293 cells were seeded into 24-well dishes and transfected the next day as described above. Plates were placed in the IncuCyte 5XS live-cell analysis system (Essen Bioscience, Ann Arbor, MI, USA) for 48 h and snapshots were taken every 30 min. Percent confluence was analyzed over time using the IncuCyte SX5 G/O/NIR Optical module software at the Bioimaging Unit, University of Haifa. Each experimental condition contained four biological repeats. Data of each condition was normalized to that value at time zero (first reading) to control for technical variability of the replicates.

### Transcriptome analyses and statistics

Total RNA was prepared from four biological replicates using the Quick-RNA MiniPrep kit (Cat. # ZR-R10554, Zymo Research). Library preparation was performed using the NEBNext Ultra RNA library Prep kit for Illumina (Cat. # E7530L, Thermo Fischer Scientific, Waltham, MA USA), according to the manufacturer’s protocol. Sequencing (single-read, 50bp) was carried out using the Illumina HiSeq 2500 at the TGC-Technion Genome Center (Technion, Haifa, Israel). Trimming of poly-A and poly-G leftovers was done using the PRINSEQ tool (https://edwards.sdsu.edu/cgi-bin/prinseq/prinseq.cgi) removing reads with more than 58% of A or G in twain with a 3’ tail filter (poly-A) of more than 8 bases. Qualified reads were later trimmed for adaptors and low-quality reads using Trimmomatic (v0.39; http://www.usadellab.org/cms/index.php?page=trimmomatic). In addition, reads were clipped to a size between 40-60 bases for optimal mapping. Gene expression levels were quantified using Htseq-count (0.6.1-py2.7) and filtered for genes covered by at least 10 reads in all samples. Differential expression was analyzed using EdgeR (3.2.4). P-values were corrected for multiple testing using the Benjamini-Hochberg method, and differential expression was considered significant for adjusted *P*-value < 0.05. The differentially expressed (DE) genes were subjected to gene-set enrichment analysis using Metascape (accessed March 2024 ^21^). Cutoff for significant enrichment was adjusted for multiple testing (Benjamini Hochberg) and the resulting q-values are reported (Tables S5 and S6).

### Statistical Analysis

Statistical analyses were done using the GraphPad Prism Software. Unless otherwise stated, statistical significance was determined by one-way ANOVA. Post-hoc analysis was performed with Dunnett’s multiple comparisons post-test when appropriate. P values < 0.05 were considered significant.

## Notes

### Competing Interest Statement

The authors have declared no competing interest.

### Summary of Updates

New data was added showing that in addition to affecting aerobic respiration, the COX-2 fragment also increases the rate of cell proliferation.

