## Supplementary material for "Activity-dependent COX-2 proteolysis generates a catalytically inactive fragment that affects aerobic respiration and proliferation": SuppFiguresAndTablesLegends.pdf (from another version of this preprint: version 1, 2024-04-11)

### Supplementary Figure Legends

**Figure S1. Gene set enrichment analysis for genes with reduced expression in CT1-expressing cells.** Top 10 most significantly enriched functional groups (as determined using Metascape) among the downregulated genes in CT1- compared to CT2-expressing cells, which showed no significant change between CT2-expressing and control cells.

### Supplementary Tables

**Table S1. Genes with significant increase in expression in CT1- compared to CT2-expressing cells.** The table contains a complete list of all the genes whose expression was significantly increased (Benjamini-Hochberg adjusted $p<0.05$) in the transcriptomic analysis.

**Table S2. Genes with significant decrease in expression in CT1- compared to CT2-expressing cells.** The table contains a complete list of all the genes whose expression was significantly decreased (Benjamini-Hochberg adjusted $p<0.05$) in the transcriptomic analysis.

**Table S3. Genes with significant increase in expression in CT1- compared to CT2-expressing cells and no change between CT2-expressing cells and untransfected cells.**
The table contains a complete list of all the genes whose expression was significantly increased (Benjamini-Hochberg adjusted $p<0.05$) in the transcriptomic analysis, but did not show differential expression (Benjamini-Hochberg adjusted $p>0.05$) between CT2-expressing cells and untransfected cells.

**Table S4. Genes with significant decrease in expression in CT1- compared to CT2-expressing cells and no change between CT2-expressing cells and untransfected cells.**
The table contains a complete list of all the genes whose expression was significantly decreased (Benjamini-Hochberg adjusted $p<0.05$) in the transcriptomic analysis, but did not show differential expression (Benjamini-Hochberg adjusted $p>0.05$) between CT2-expressing cells and untransfected cells.

**Table S5. Gene sets showing significant enrichment among genes upregulated in CT1- compared to CT2-expressing cells and not changing between CT2-expressing cells and untransfected cells.**
Metascape was used to perform the analysis.

**Table S6. Gene sets showing significant enrichment among genes downregulated in CT1- compared to CT2-expressing cells and not changing between CT2-expressing cells and untransfected cells.** Metascape was used to perform the analysis.
