## Supplementary figures and images for "Activity-dependent COX-2 proteolysis generates a catalytically inactive fragment that affects aerobic respiration and proliferation" (from another version of this preprint: version 1, 2024-04-11)

### Figure S1.tiff

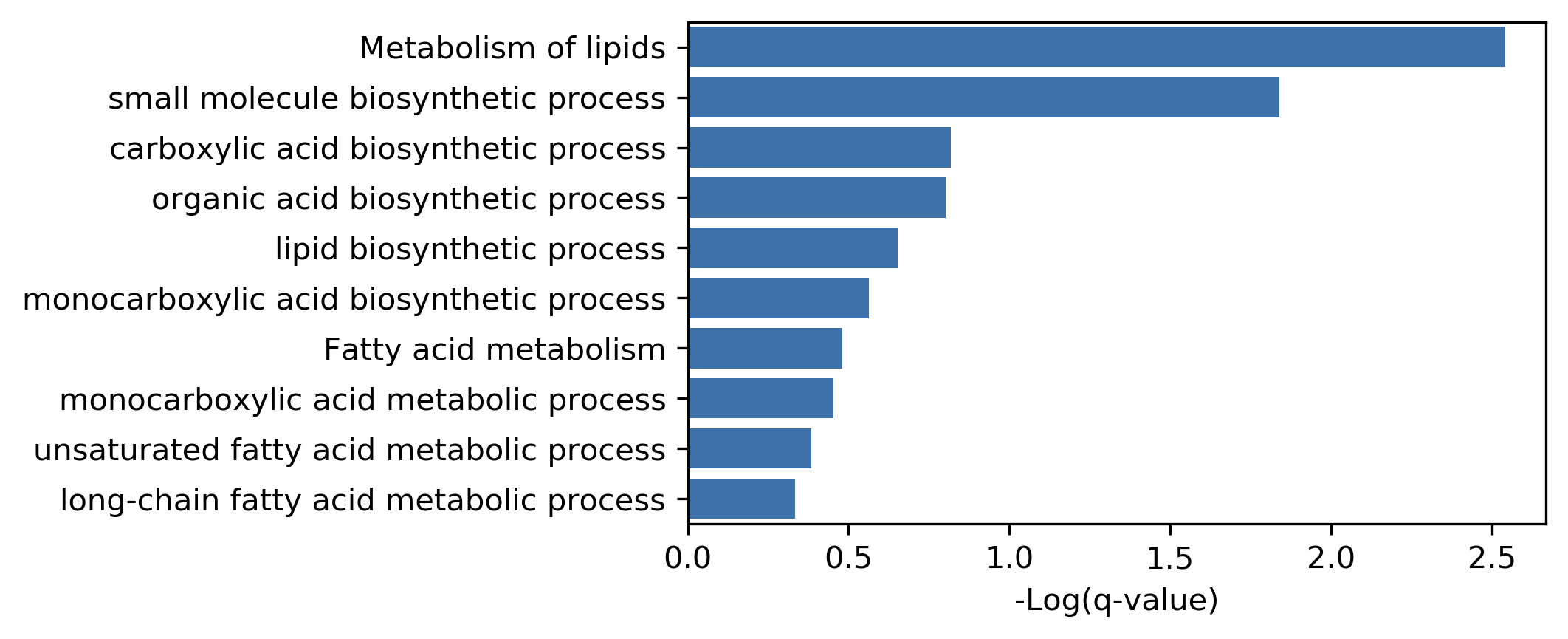
